# The liver compensates for zonal loss of glucose 6-phosphatase to prevent glycogen storage disease

**DOI:** 10.64898/2026.07.26.740786

**Authors:** Roger Liang, Ling Cai, Phong T. Nguyen, Sherwin Kelekar, Maggie B. Cervantes, Trevor Tippetts, Eric Chen, Roberto Ribas, Alison Ryan, Janice Chou, Ramon Sun, Hao Zhu, Ralph J. DeBerardinis

## Abstract

The liver is organized into spatial zones with distinct metabolic roles, but how this architecture contributes to disease remains unclear. Glucose production is concentrated in periportal hepatocytes and depends on G6PC1; loss of this enzyme causes glycogen storage disease type Ia (GSD1a). We tested whether G6PC1 loss in specific zones, including its primary periportal location, is sufficient to cause disease. Unexpectedly, loss of *G6pc1* in any single zone caused local glycogen accumulation but did not produce the systemic metabolic abnormalities or liver tumors seen after whole-liver deletion. Instead, disease developed only after near-complete loss of *G6pc1* across the entire liver. These findings show that organ-wide compensation preserves metabolic homeostasis and suppresses tumorigenesis despite localized disruption.

**Teaser:** Spatial *G6pc1* loss causes local glycogen storage without systemic metabolic disease.

## Introduction

The liver is a spatially organized tissue in which hepatocytes adopt distinct metabolic programs within lobules depending on their position along the porto-central axis. This organization, termed metabolic zonation, is regulated by gene expression programs dictated by gradients of oxygen, nutrients, and hormones as blood flows from the portal vein through sinusoids into the central vein (*1*). Periportal (zone 1) hepatocytes, located near the portal triad, are enriched for enzymes involved in gluconeogenesis, glycogenolysis, β-oxidation, and ureagenesis; midlobular (zone 2) hepatocytes are responsible for repopulating the liver; and pericentral (zone 3) hepatocytes near the central vein specialize in glycolysis, lipogenesis, and xenobiotic metabolism (*2–4*). This division of labor is thought to allow the liver to sustain systemic homeostasis during fluctuating metabolic demands, despite regional heterogeneity of substrate availability among hepatocytes (*5*).

Despite detailed transcriptomic and proteomic characterization of zonation, its functional significance and robustness to metabolic perturbation remain poorly understood (*6*). Most studies in mice employ whole-body or whole-liver perturbations, obscuring zone-specific contributions. Recent work using zonal gene manipulation in insulin signaling and oncogenic pathways has demonstrated that spatially restricted perturbations lead to unexpected metabolic and malignant phenotypes (*7-8*). However, the genes in these studies have broad, pleiotropic functions, complicating mechanistic interpretation. Thus, a key unanswered question is whether zonal inhibition of a single, well-defined metabolic reaction is sufficient to produce systemic disease or to alter the behavior of neighboring wild-type (WT) hepatocytes.

Gluconeogenesis and glycogenolysis, which maintain glucose homeostasis during fasting, converge on glucose 6-phosphatase catalytic subunit 1 (G6PC1)-mediated glucose-6-phosphate (G6P) hydrolysis. Loss-of-function mutations in *G6PC1* cause glycogen storage disease type 1a (GSD1a), an autosomal recessive inborn error of metabolism characterized by systemic metabolic anomalies (fasting hypoglycemia, lactic acidosis, hyperuricemia, and hypertriglyceridemia) and a high incidence of hepatic adenomas that can progress to hepatocellular carcinoma (*9–11*). Whole-liver knockout (KO) of *G6pc1* in mice recapitulates these metabolic defects and predisposes the liver to tumorigenesis. Liver cancers in this model are thought to arise from cell-autonomous signaling effects of glycogen accumulation (*12–14*). In mice and humans, gluconeogenic and glycogenolytic activity are highest in zone 1 hepatocytes, providing a clear, testable prediction. If zonation is functionally essential for glucose homeostasis, then deleting *G6pc1* in zone 1 should reproduce the metabolic and oncogenic manifestations of whole-liver deficiency (*15–16*). To address this question, we used tamoxifen-inducible CreER drivers to delete *G6pc1* in the entire liver or in spatially-defined hepatocyte subpopulations. This allowed us to directly test whether spatially confined inhibition of gluconeogenesis and glycogenolysis is sufficient to generate the systemic features of GSD1a. By functionally perturbing a single, well-defined metabolic reaction within specific zones, we could experimentally define how zonation shapes metabolism and tumor susceptibility.

## Results

### G6PC1 and glycogen metabolism are enriched in periportal hepatocytes

We performed spatial transcriptomics on mouse livers using the Xenium platform with a custom 500-gene panel enriched for transcripts related to liver metabolic zonation. We observed high *G6pc1* mRNA levels in zone 1 hepatocytes that co-express the periportal marker *Gls2* (fig. S1A). To define the spatial distribution of G6PC1 protein expression, we examined published spatial proteomics data from Ben-Moshe et al., which demonstrated that glycolysis/gluconeogenesis was among the most strongly zonated pathways in the liver, with enrichment in periportal hepatocytes (fig. S1B) (*6*). Gradients for individual gluconeogenic enzymes increased from the central vein toward the portal vein, mirroring the distribution of glycogen metabolism enzymes, such as glycogen synthase (GYS2) and glycogen branching enzyme (GBE1) (fig. S1C-E). Together, these data confirm the periportal enrichment of *G6pc1* and establish a spatial baseline for interpreting the physiological consequences of its zone-specific deletion.

### Zone-specific *G6pc1* deletion does not mimic GSD1a

Wei et al. previously developed zone-specific CreER (*zone-CreER*) mice to enable selective genetic manipulation of hepatocytes from each metabolic zone (*17-18*). To generate zone-specific *G6pc1* KO models, we crossed *G6pc1^fl/fl^* mice with CreER drivers targeting periportal (*Gls2-CreER*; zone 1), midlobular (*Igfbp2-CreER*; zone 2), or pericentral (*Cyp1a2-CreER*; zone 3) hepatocytes. Mice with the *G6pc1^fl/fl^* lacking CreER were used as a negative control, and *Alb-CreER;G6pc1^fl/fl^* mice were used as a positive control for whole-liver *G6pc1* deletion. The latter model has been shown to induce the physiological defects of GSD1a, including liver glycogen accumulation, hepatomegaly, fasting hypoglycemia, and elevated serum lactate (*12*).

We first tested different tamoxifen regimens to induce *G6pc1* deletion across all CreER models within the same experiment. Mice carrying heterozygous CreER alleles were assigned to either five consecutive daily IP tamoxifen injections or tamoxifen-containing chow administered ad libitum for 2.5 weeks (fig. S2A). This side-by-side comparison allowed us to directly evaluate whether tamoxifen delivery affected recombination efficiency across the whole-liver and zone-specific *G6pc1* KO models. For immunofluorescence analysis, GS staining was used to mark the pericentral zone 3 region and orient the lobule, whereas loss of G6PC1 protein signal was used as the readout for *G6pc1* deletion (fig. S2B-F). IP tamoxifen efficiently deleted *G6pc1* in the *Alb-CreER* whole-liver KO model and produced robust recombination in the *Cyp1a2-CreER* zone 3 KO model (fig. S2B-C, and F). However, the same IP regimen produced undetectable recombination in the *Gls2-CreER* zone 1 or Igfbp2-CreER zone 2 models, as indicated by preserved G6PC1 expression and minimal glycogen-associated vacuolization (fig. S2D-E). In the same experimental cohort, tamoxifen chow produced zone-restricted histologic changes on H&E and corresponding loss of G6PC1 protein in each CreER model (fig. S2C-F). We quantified G6PC1-negative cells as an image-based estimate of *G6pc1* recombination efficiency in the whole liver. The analysis showed that tamoxifen chow significantly increased the percentage of G6PC1-negative cells in the zone 1 and zone 2 KO models, while maintaining efficient recombination in the whole-liver and zone 3 KO models (fig. S2G-K). After tamoxifen chow, each CreER model contained more G6PC1-negative cells than no-CreER controls (fig. S2L). Thus, within a single side-by-side induction experiment, tamoxifen chow provided the most reliable regimen for producing measurable *G6pc1* deletion across all zone-specific CreER models.

Having established tamoxifen chow as a reliable induction regimen, we next used this approach to phenotype an expanded cohort of whole-liver and zone-specific *G6pc1* KO mice. This cohort included both heterozygous and homozygous *Gls2-CreER* and *Cyp1a2-CreER* mice. G6PC1 signal was reduced in the expected spatial regions of each zone-specific model and broadly eliminated in the whole-liver KO model (Fig. 1A–E). Compared with no-CreER controls, each *G6pc1* KO model contained significantly more G6PC1-negative cells (Fig. 1F). The zone 1 and zone 3 KO models contained an average of 46% and 47% G6PC1-negative cells, respectively. Qualitative and quantitative assessment of these models suggested that G6PC1 loss extended beyond the most restricted periportal or pericentral regions and into adjacent midlobular hepatocytes.

**Figure 1.**
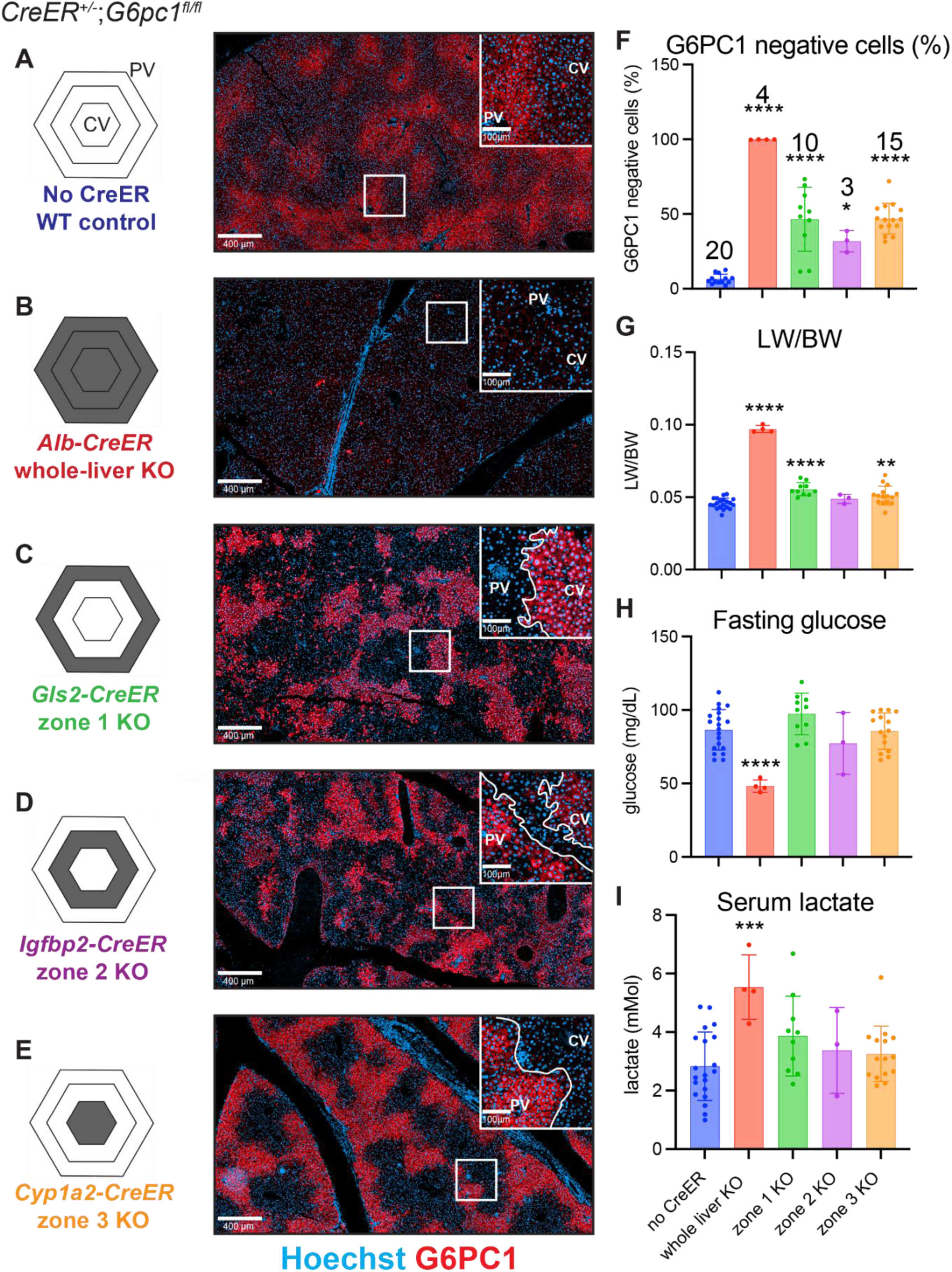
Zonal G6pc1 deletion does not cause systemic GSD1a-like metabolic dysfunction. (**A–E**) Representative G6PC1 immunofluorescence images from no-CreER control, *Alb-CreER* whole-liver KO, *Gls2-CreER* zone 1 KO, *Igfbp2-CreER* zone 2 KO, and *Cyp1a2-CreER* zone 3 KO livers 1 month after tamoxifen chow induction. Insets show higher-magnification views with portal vein (PV) and central vein (CV) regions indicated. White outlined regions indicate areas of G6PC1 loss used to identify recombined/KO regions. (**F**) Quantification of G6PC1-negative cells as a percentage of all cells in the liver. (**G**) Liver weight/body weight ratio. (**H**) Fasting blood glucose. (**I**) Serum lactate. Zone 1 and zone 3 KO groups include heterozygous and homozygous CreER mice; stratified zone 1 analysis is shown in Fig. S3. Data represent mean ± SD. One-way ANOVA followed by Tukey’s multiple comparisons test was used to compare each group with the no-CreER control. Statistical significance was denoted as follows: *, P < 0.05; **, P < 0.01; ***, P < 0.001; ****, P < 0.0001.

Despite substantial zonal recombination, zone 1 and zone 3 KO mice displayed only a small increase in liver weight/body weight ratio (LW/BW) compared with no-CreER controls (Fig. 1G). More importantly, *G6pc1* deletion from any single metabolic zone failed to disrupt systemic glucose homeostasis. After a 9-hour fast, zone 1, zone 2, and zone 3 KO mice maintained normal blood glucose and serum lactate, whereas whole-liver KO mice developed fasting hypoglycemia and hyperlactatemia, as expected (Fig. 1H–I). Thus, zonal *G6pc1* deletion produced measurable local liver effects but did not recapitulate the systemic metabolic abnormalities of whole-liver *G6pc1* deficiency.

Because zone 1 is the major site of G6PC1-dependent glucose production, we performed a stratified analysis of zone 1 mice from the same expanded cohort to determine whether increasing zone 1 recombination induced systemic GSD1a-like phenotypes. Mice with homozygous *Gls2-CreER^+/+^*had increased *G6pc1* recombination compared with heterozygous *Gls2-CreER^+/-^*mice (fig. S3A-B). However, increasing recombination did not result in significant changes in LW/BW, fasting glucose, or serum lactate (fig. S3C-E). Thus, increasing zonal *G6pc1* deletion strengthened local liver phenotypes in some settings but remained insufficient to produce systemic GSD1a-like metabolic dysfunction. Because homozygous *CreER* alleles increased recombination without altering the systemic phenotype, subsequent zonal KO cohorts included heterozygous and homozygous CreER mice.

Because *G6pc1* deletion from any single metabolic zone did not produce systemic disease, we next asked how much total hepatic *G6pc1* loss is required to disrupt glucose homeostasis. To generate graded, zonally agnostic recombination across the liver, we administered AAV8-TBG-Cre to *G6pc1^f/l/fl^* mice at low (2×10*^9^* vector genomes: vg), medium (1×10*^10^* vg), and high (5×10*^10^* vg) doses (Fig. 2A). Mice had 31%, 89%, and 97% G6PC1*-* negative cells, respectively (Fig. 2B). Mice at the medium dose displayed hepatomegaly without hypoglycemia, and only mice at the highest dose developed both hepatomegaly and fasting hypoglycemia (Fig. 2C-D). These findings supported the conclusion that near-complete, rather than spatially localized, loss of G6PC1 is required to cause GSD1a-like systemic abnormalities.

**Figure 2.**
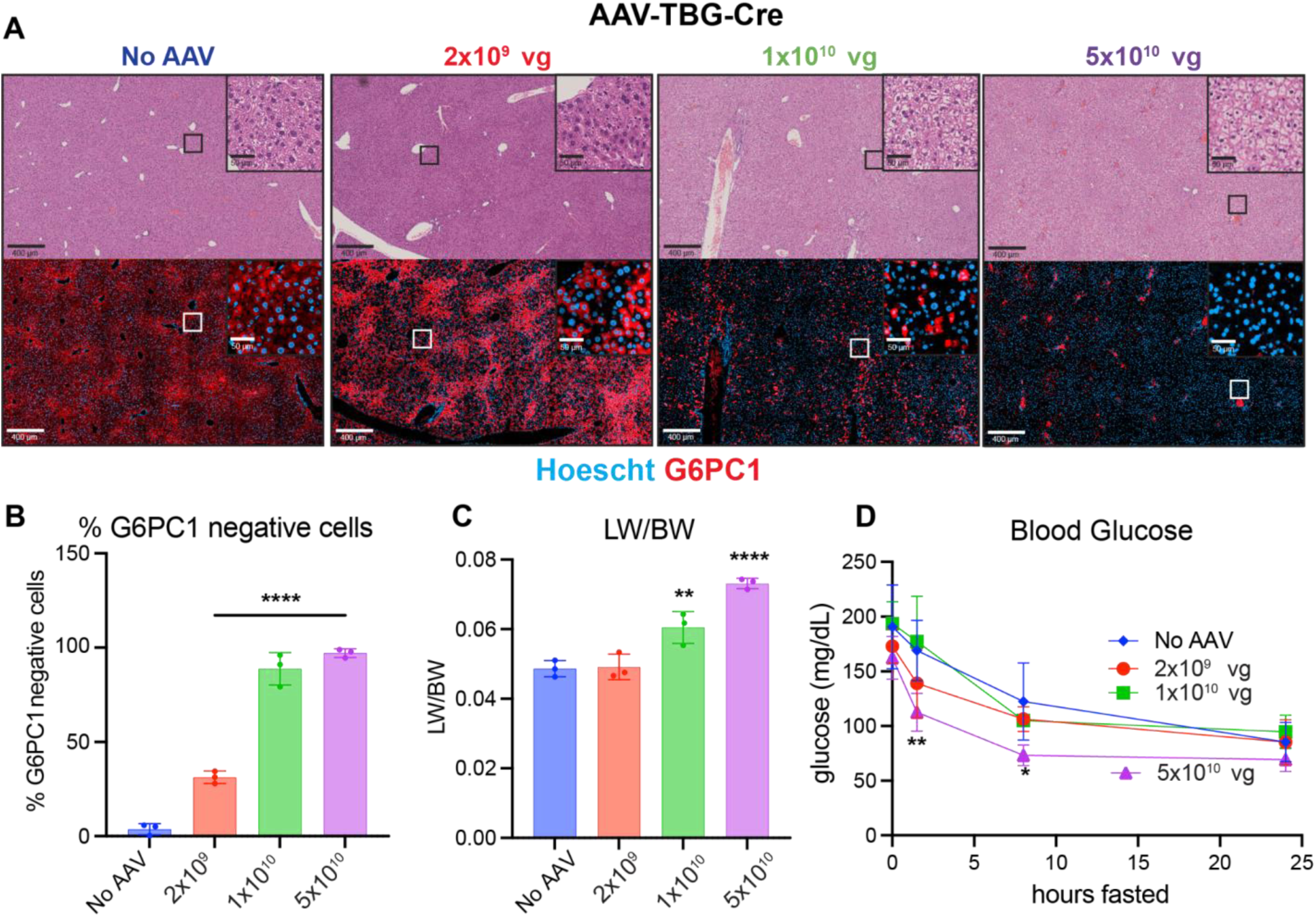
Near-complete liver-wide *G6pc1* deletion is required for fasting hypoglycemia. (**A**) Representative G6PC1 IF in mice treated with escalating doses of AAV8-TBG-Cre. (**B**) Quantification of *G6pc1* deletion, measured as the percentage of G6PC1-negative cells, after low (2×10^9^), medium (1×10^10^), or high (5×10^10^) dose of AAV8-TBG-Cre (**C**) LW/BW ratios, and (**D**) fasting glucose in mice given escalating doses of AAV-TBG-Cre. Data represent mean ± SD. One-way ANOVA analyses were followed by Tukey’s multiple comparisons test comparing each group to the no-AAV control in (**B-C**). Two-way ANOVA analyses were followed by Fisher’s least significant difference (LSD) test for multiple comparisons in (**D**). Statistical significance was denoted as follows: *, P < 0.05; **, P < 0.01; ***, P < 0.001, ****, P < 0.0001. Vg, viral genomes.

### Spatial metabolomics of zonal *G6pc1* KOs reveals zone-specific glycogen accumulation

After establishing that systemic metabolic dysfunction required near-complete hepatic G6PC1 loss, we next asked whether zonal *G6pc1* deletion produced local metabolic abnormalities within the liver. We performed Matrix-Assisted Laser Desorption/Ionization Time-of-Flight (MALDI-ToF) mass spectrometry imaging on liver sections from no-CreER control, *Gls2-CreER^+/+^* zone 1 KO, *Cyp1a2-CreER^+/+^* zone 3 KO, and whole-liver KO mice. We focused on the zone 1 and zone 3 models because they represent the strongest metabolic contrast across the porto-central axis, whereas zone 2 is intermediate and showed the same absence of systemic metabolic dysfunction as the other zonal KOs.

MALDI-ToF imaging enabled visualization of the relative spatial distributions of glycogen species, aqueous small-molecule metabolites, and lipids in liver sections (Fig. 3A-B). Because hepatic glycogen metabolism and G6PC1-dependent glucose production are spatially zonated, we expected control livers to display heterogeneous distributions of glycogen-associated metabolites. In zonal *G6pc1* knockout livers, we expected glycogen and the G6PC1 substrate glucose-6-phosphate (G6P) to accumulate preferentially in regions containing *G6pc1* KO hepatocytes, whereas whole-liver *G6pc1* KO was expected to produce broader accumulation across the lobule. Consistent with this prediction, a representative glycogen-associated feature, m/z 1175, showed heterogeneous spatial abundance in no-CreER control livers and localized enrichment in the zone 1 and zone 3 KO livers (Fig. 3A). In contrast, whole-liver KO sections showed more homogeneous accumulation of this glycogen-associated feature across the tissue, consistent with glycogen storage throughout the liver (Fig. 3A). G6P was also heterogeneously distributed in no-CreER control and zonal knockout livers, whereas whole-liver knockout sections showed more widespread and uniform G6P accumulation (Fig. 3B). Together, these spatial metabolomics data indicate that zonal *G6pc1* deletion produces localized metabolic abnormalities, while whole-liver deletion causes more global accumulation of glycogen-associated metabolites and G6P.

**Figure 3.**
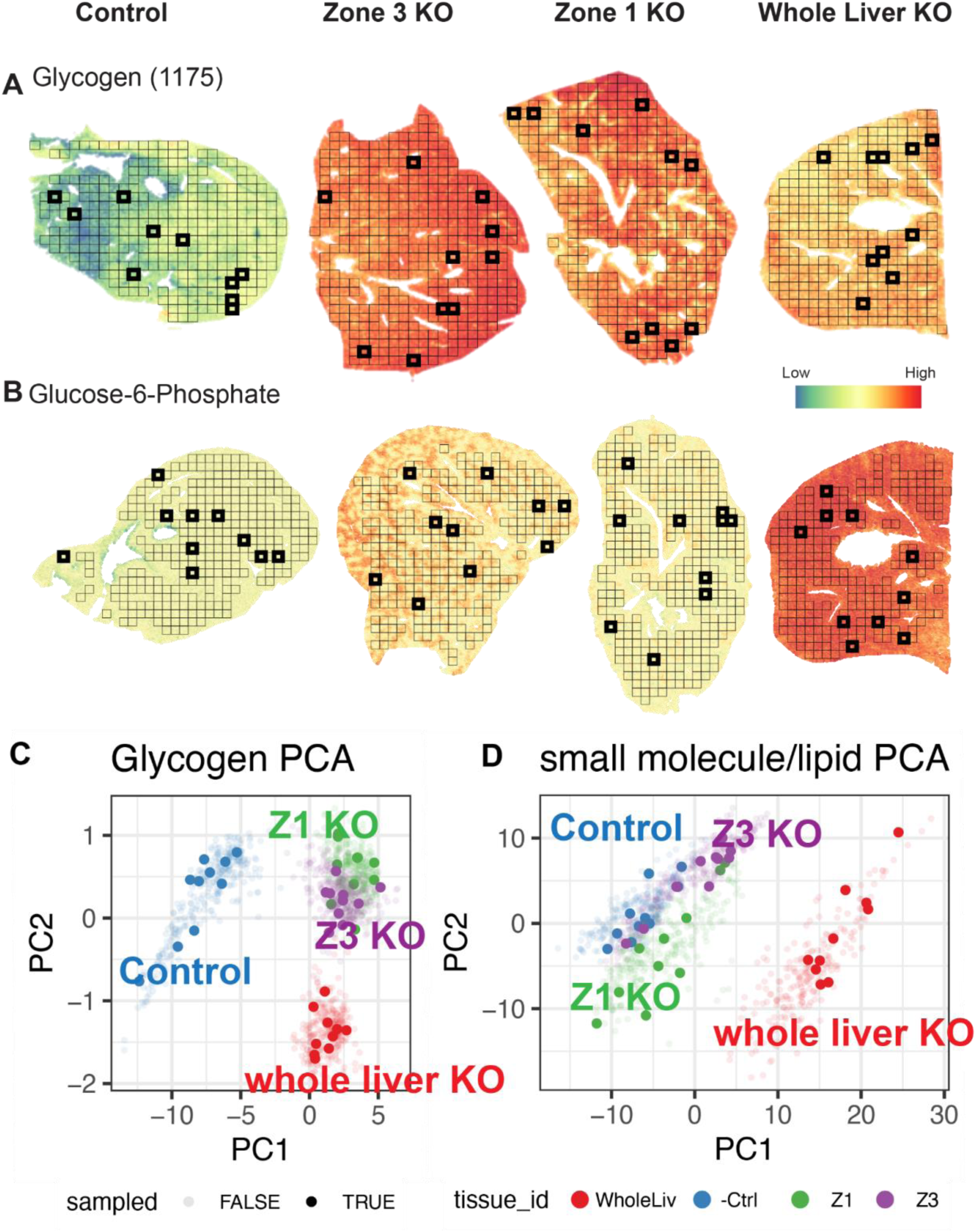
Spatial metabolomics reveals localized glycogen and G6P accumulation in zonal *G6pc1* KO livers. MALDI-ToF images showing the relative spatial distribution of (**A**) a representative glycogen-associated feature, m/z 1175 and (**B**) glucose-6-phosphate (G6P). Metabolite abundance is visualized from a continuous gradient from low (1st percentile) to high (99th percentile) with global scaling. Black bins were used to generate PCA plots (**C-D**). Tile-level analyses were used for descriptive spatial comparison and not for genotype-level statistical inference.

To compare these spatial patterns across sections, we performed tile-level analyses of glycogen-associated features and small-molecule/lipid features. PCA analysis of glycogen-associated features showed that zone 1 and zone 3 KO sections shared similar glycogen profiles that were distinct from both no-CreER controls and whole-liver KOs (Fig. 3C). In contrast, PCA of small-molecule and lipid features showed that no-CreER control and zonal KO sections clustered more closely together, whereas whole-liver KO sections separated from the other groups (Fig. 3D). Heatmap analysis of the same tile-level datasets supported these broad differences in glycogen-associated, small-molecule, and lipid features across control, zonal KO, and whole-liver KO sections (fig. S4). Because MALDI tiles are spatially correlated measurements from individual tissue sections rather than independent biological replicates, these analyses were used as descriptive spatial support rather than genotype-level statistical inference. Together, MALDI-ToF imaging showed that zonal *G6pc1* deletion produced localized glycogen and G6P abnormalities without the pronounced metabolic remodeling observed after whole-liver *G6pc1* deletion.

### A glycogen sensor reveals reciprocal glycogen redistribution after zonal *G6pc1* deletion

MALDI-ToF imaging suggested that zonal *G6pc1* KO produced localized glycogen abnormalities that were distinct from the control or whole-liver KO models. However, the spatial resolution of MALDI-ToF was insufficient to determine whether glycogen accumulated specifically in G6PC1-deficient hepatocytes. We therefore sought to visualize glycogen at cellular resolution and multiplex this signal with G6PC1 IF. H&E staining can suggest glycogen accumulation through cytoplasmic clearing and vacuolization, but periodic acid– Schiff (PAS) staining is the standard histological method for detecting liver glycogen (*19*). However, PAS staining is not readily multiplexed with G6PC1 IF on the same tissue section. To overcome this limitation, we developed and purified a fluorescent glycogen sensor that can be multiplexed with antibodies. The glycogen sensor conjugated an mCherry construct with the glycogen-binding CBM20 domain of Starch-binding domain-containing protein 1 (STBD1), which binds to glycogen to facilitate glycophagy (*20-21*).

We first tested whether the glycogen sensor detected expected hepatic glycogen patterns. In control livers, sensor signal was enriched in periportal hepatocytes, consistent with the known spatial distribution of hepatic glycogen (*15-16*, *22*)(Fig. 4A). In whole-liver *G6pc1* KO sections, glycogen was detected across the lobule as expected (Fig. 4A). To confirm that the sensor signal was glycogen-dependent, we pretreated slides with α-amylase, which digests glycogen. Doing so abolished sensor signal in both the control and whole-liver KO sections, supporting the specificity of the sensor for glycogen (Fig. 4A).

**Figure 4.**
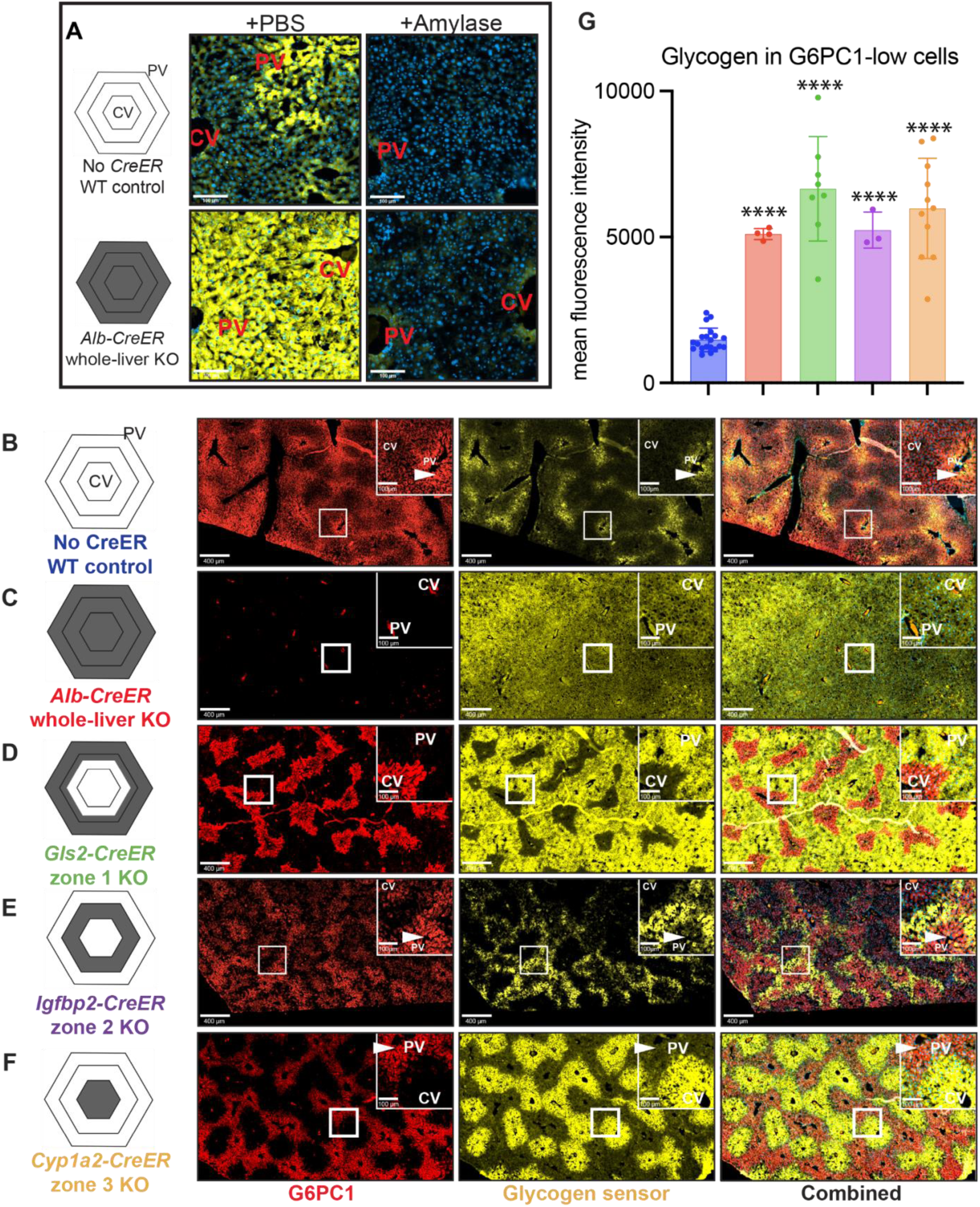
Zonal *G6pc1* KO hepatocytes accumulate glycogen at levels comparable to whole-liver KO. (**A**) A glycogen sensor (mCherry-6xHis-FLAG-CBM20) was purified and tested on WT and whole-liver KO frozen liver sections with or without α-amylase pretreatment. α-amylase digestion abolished sensor signal, supporting glycogen-dependent staining. (**B-F**) Representative multiplex immunofluorescence images showing G6PC1 and glycogen sensor staining in no-CreER control, whole-liver KO, and zone-specific *G6pc1* KO liver sections from fasted mice. Arrows indicate periportal regions with high glycogen in control liver and relative glycogen depletion in corresponding regions of zone 2 and 3 KO livers. (**G**) Quantification of glycogen sensor intensity in G6PC1-low hepatocytes across control, whole-liver KO, and zonal KO sections. Data represent mean ± SD. One-way ANOVA followed by Tukey’s multiple comparisons test was used to compare each group. Zonal KO groups were significantly higher than no-CreER control and were not significantly different from whole-liver KO. Statistical significance was denoted as follows: *, P < 0.05; **, P < 0.01; ***, P < 0.001; ****, P < 0.0001.

We then applied both the glycogen sensor and G6PC1 IF to liver sections from the entire series of *G6pc1* KO models (Fig. 4B-F). In no-CreER controls, glycogen signal was highest in periportal hepatocytes, which also expressed high levels of G6PC1 (Fig. 4B). In whole-liver KOs, glycogen signal and G6PC1 loss were broadly distributed (Fig. 4C). In zone 1, zone 2, and zone 3 KO livers, glycogen accumulated in the corresponding regions of G6PC1 loss (Fig. 4D–F). Accumulation in the targeted zones seemed to affect glycogen distribution in remaining wild-type hepatocytes. For example, after zone 2 and zone 3 deletion, glycogen accumulation in midlobular or pericentral regions was accompanied by reduced glycogen signal in periportal regions that normally contain abundant glycogen (Fig. 4B, E, and F).

To quantify glycogen accumulation in recombined hepatocytes, we segmented cells and classified hepatocytes as G6PC1-low based on cytoplasmic G6PC1 IF intensity. We used the term G6PC1-low, rather than G6PC1-negative, for this analysis because the threshold was chosen to capture the physiologic low-G6PC1 hepatocyte population present in no-CreER control livers, allowing this control population to serve as a comparator across genotypes. G6PC1-low hepatocytes from each *G6pc1* KO model contained much more glycogen than G6PC1-low hepatocytes from no-CreER control livers (Fig. 4G). Moreover, glycogen abundance in G6PC1-low hepatocytes from zonal KO livers was similar to whole-liver KO hepatocytes, indicating that zonal KO cells accumulated glycogen to levels comparable to whole-liver *G6pc1* deficiency (Fig. 4G). Thus, localized *G6pc1* deletion produced a robust cell-level glycogen-storage phenotype, even though these mice did not develop systemic GSD1a-like metabolic dysfunction.

### Zonal *G6pc1* KO hepatocytes persist long term

Glycogen accumulation has been proposed to confer a selective advantage to premalignant hepatocytes through inhibition of the Hippo pathway (*27*). Thus, we tested whether G6PC1-negative, glycogen-loaded hepatocytes would be preferentially retained or expand over time after zonal *G6pc1* deletion. We aged zonal KO mice for 12 months after tamoxifen induction and quantified the persistence of G6PC1-negative cells by IF (Fig. 5A-E). For the zone 1 analysis, we used *Gls2-CreER^+/−^* mice to preserve periportal restriction of the recombined population.

**Figure 5.**
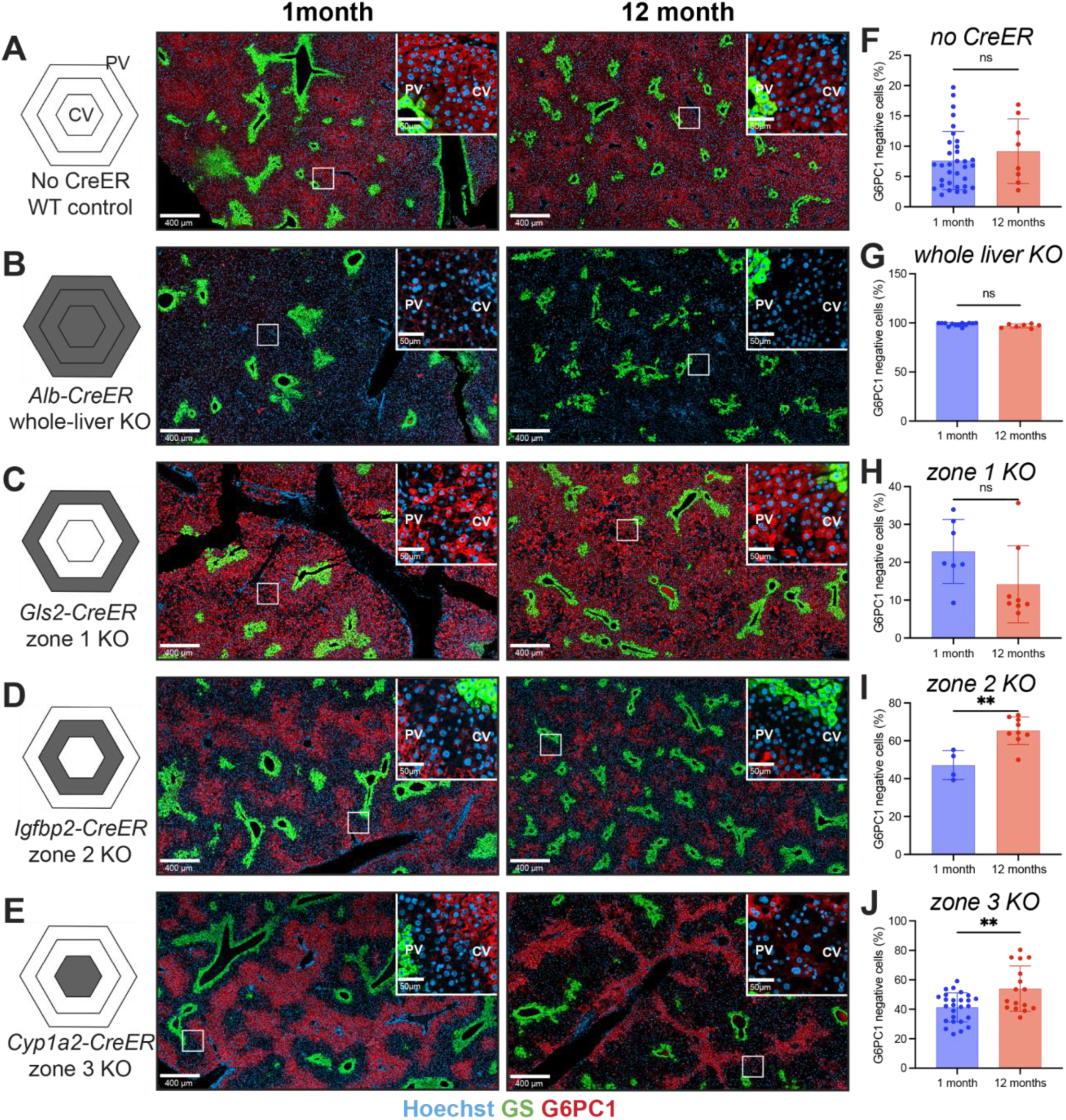
G6PC1-negative hepatocytes persist long term after zonal *G6pc1* deletion. Representative G6PC1 IF performed 1 and 12 months after tamoxifen administration to investigate persistence of G6PC1-negative cells in mice with (**A**) no creER, (**B**) whole-liver KO, (**C**) zone 1 KO, (**D**) zone 2 KO, and (**E**) zone 3 KO. GS staining marks pericentral hepatocytes and was used to orient the lobule. For the zone 1 persistence analysis, Gls2-CreER^+/−^ mice were used to preserve periportal zone restriction. (**F–J**) Quantification of G6PC1-negative cells at 1 and 12 months after tamoxifen induction in the corresponding models. Data represent mean ± SD. Unpaired two-tailed t tests were used to compare 1-month and 12-month groups within each model. Statistical significance was denoted as follows: *, P < 0.05; **, P < 0.01; ***, P < 0.001; ****, P < 0.0001.

No-CreER control mice displayed few G6PC1-negative cells, whereas whole-liver G6pc1 KO mice retained widespread G6PC1 loss (Fig. 5A and B). G6PC1-negative cells remained detectable in all zonal KO models at 12 months after induction (Fig. 5C–E). Between 1 and 12 months after tamoxifen, the fraction of G6PC1-negative cells was unchanged in no-CreER and whole-liver KO controls, trended lower in zone 1 KOs, and increased in zone 2 and zone 3 KOs (Fig. 5F–J). These zone-dependent changes paralleled the lineage behavior previously reported for tomato-labeled hepatocytes under homeostatic conditions, in which zone 1-labeled cells decline while zone 2– and zone 3-labeled cells expand over time (*17-18*). Thus, *G6pc1*-deficient hepatocytes persisted over 12 months, but *G6pc1* loss and glycogen accumulation in these models did not override the baseline zone-dependent behavior of hepatocyte populations.

### Zonal *G6pc1* deletion does not recapitulate the whole-liver KO tumor phenotype

Some inborn errors of metabolism, including GSD1a, are associated with an elevated risk of liver tumorigenesis. GSD1a is unusual in that tumors arise in the absence of cirrhosis, suggesting that the tumors result from the primary metabolic disturbance rather than chronic liver injury (*23-24*). Although local glycogen accumulation did not confer an obvious selective advantage to zonal KO hepatocytes, it was possible that the glycogen accumulation would still result in increased tumorigenesis, as premalignant cell proliferation has previously been uncoupled from tumorigenesis (*8*). Thus, we tested the impact of regional *G6pc1* deficiency on hepatic tumor formation.

As a positive control, we first confirmed that all whole-liver *G6pc1* KO mice developed gross liver tumors 12 months after *G6pc1* deletion (fig. S5A) (*12*). These mice did not have increased serum ALT, consistent with tumor formation in the absence of liver injury (fig. S5B). To further characterize this tumor phenotype, we performed whole-exome sequencing on 10 tumors from whole-liver KO mice. These tumors had a low mutational burden, with a median of 0.13 somatic variants per megabase, compared to the much higher mutational burdens in human hepatocellular carcinomas in the LIHC database and diethylnitrosamine-induced mouse liver tumors (fig. S5C). We did not identify recurrent mutations in known liver cancer driver genes among the sequenced tumors (*25*, *26*). Together, these findings support the use of whole-liver *G6pc1* KO mice as a metabolically driven positive-control model of GSD1a-associated liver tumorigenesis.

Despite long-term persistence of glycogen-loaded G6PC1-negative cells, zonal KO mice did not recapitulate the macroscopic tumor phenotype of whole-liver *G6pc1* deletion. After long-term aging, whole-liver KO mice developed overt liver tumors, whereas zonal KO livers did not develop comparable tumor burden on gross exam (Fig. 6A–E). H&E staining showed preserved overall tissue architecture in no-CreER control livers, whereas whole-liver KO tumors displayed diffuse cytoplasmic clearing, consistent with the glycogen-storage phenotype associated with *G6pc1* loss (Fig. 6F–G). In contrast, H&E staining of zonal KO livers showed zone-restricted cytoplasmic changes without the extensive tumor burden observed in whole-liver KO livers (Fig. 6H– J). Consistent with the gross and histologic findings, whole-liver KO mice developed marked hepatomegaly, whereas zonal KO mice maintained substantially lower LW/BW ratios across 10-, 12-, and 15-month cohorts (Fig. 6K–M). Zonal KO mice also maintained fasting blood glucose, whereas whole-liver KO mice maintained systemic GSD1a-like metabolic dysfunction, indicating that zonal KO mice did not develop systemic GSD1a-like disease even after long-term aging (Fig. 6N–P). These results indicate that localized *G6pc1* loss and chronic glycogen accumulation are insufficient to recapitulate the penetrant macroscopic tumor and systemic metabolic phenotypes produced by whole-liver *G6pc1* deletion.

**Figure 6.**
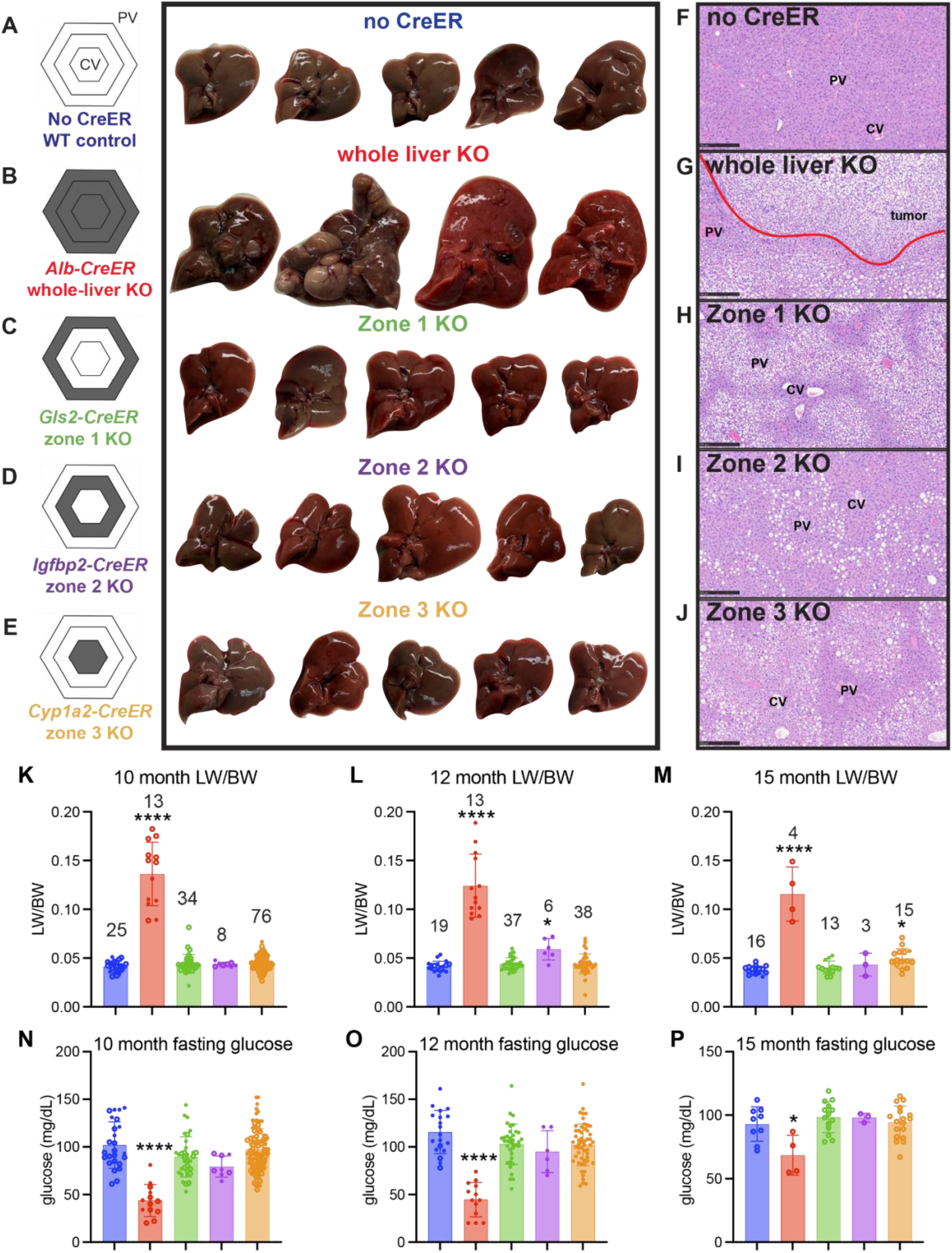
Zonal *G6pc1* deletion fails to recapitulate whole-liver KO tumor burden. Representative gross liver images from (**A**) no-CreER control, (**B**) whole-liver KO, (**C**) zone 1 KO, (**D**) zone 2 KO, and (**E**) zone 3 KO mice 12 months after tamoxifen chow induction. Whole-liver KO mice developed liver tumors, whereas zonal KO mice did not develop comparable macroscopic tumor burden. (**F–J**) Representative H&E-stained liver sections from the corresponding models. Whole-liver KO livers contained tumors with diffuse cytoplasmic clearing, outlined in red. Zonal KO livers generally showed preserved tissue architecture with zone-restricted cytoplasmic changes. (**K–M**) Liver weight/body weight ratio and (**N–P**) fasting blood glucose in mice analyzed 10, 12, or 15 months after tamoxifen induction. Numbers above bars indicate the number of mice per group. Data represent mean ± SD. One-way ANOVA followed by Tukey’s multiple comparisons test was used to compare each group with the no-CreER control. Statistical significance was denoted as follows: *, P < 0.05; **, P < 0.01; ***, P < 0.001; ****, P < 0.0001.

## Discussion

Our findings reveal that the liver can buffer spatially restricted loss of a key glucose-producing enzyme. Although G6PC1 is strongly enriched in periportal hepatocytes and is essential for gluconeogenesis and glycogenolysis, *G6pc1* deletion from any single metabolic zone caused local glycogen accumulation without fasting hypoglycemia, lactate elevation, or overt liver tumor formation. Consistent with this buffering capacity, fasting hypoglycemia emerged only after *G6pc1* was deleted from nearly all hepatocytes, indicating a steep tissue-level threshold for metabolic dysfunction in this model. These findings complement gene therapy studies in germline and liver-specific *G6pc1*^-/-^ mice, which show that complete restoration of hepatic G6Pase-α activity is not required for therapeutic benefit and that low residual activity can maintain glucose homeostasis and prevent liver tumorigenesis (*28–30*). Our results extend this principle by suggesting that therapeutic efficacy may depend more on preserving sufficient liver-wide G6PC1 activity than on restoring expression specifically to its native periportal zone. This interpretation is consistent with recent evidence that gluconeogenesis can be redistributed across the lobule during metabolic transitions (*31*). We emphasize that in our models, G6PC1 is expressed in the kidney, another organ capable of glycogenolysis and gluconeogenesis that may contribute to systemic metabolic homeostasis. Nevertheless, the data demonstrate that extensive periportal G6PC1 expression is dispensable for glucose homeostasis in mice.

The ability of zonal *G6pc1* KO mice to maintain systemic glucose homeostasis was associated with glycogen remodeling across the lobule. Using our engineered glycogen sensor, we found that glycogen accumulated in regions of G6PC1 loss, demonstrating that zonal *G6pc1* deletion produced a local glycogen-storage phenotype at cellular resolution and that this occurred beyond zone 1. After zone 2 and zone 3 deletion, glycogen accumulation in the targeted region was accompanied by relative depletion of glycogen in periportal regions where glycogen normally accumulates. Thus, localized *G6pc1* loss does not simply add glycogen to the targeted region; instead, it affects glycogen distribution in remaining wild-type hepatocytes. This pattern is consistent with tissue-level buffering by intact hepatocytes, although the mechanism underlying this response remains unknown.

Our results also suggest a need to revise how glycogen storage impacts GSD1a-associated liver tumorigenesis. We initially hypothesized that zonal *G6pc1* deletion might produce a spatial bias in tumorigenesis because prior work demonstrated that metabolic zonation is a determinant for *Ctnnb1-*driven liver tumor initiation and that hepatic glycogen overload can promote tumor initiation through Hippo-pathway suppression (*8*, *27*). Although G6PC1-negative cells in the zonal KO accumulated as much glycogen as those in the whole-liver KO, zonal *G6pc1* deletion did not induce tumorigenesis even 15 months after tamoxifen. Thus, localized glycogen accumulation was not sufficient for overt liver tumor formation in this model. Tumor risk is also unlikely to be determined solely by total residual hepatic G6PC1 activity, because low residual activity can prevent tumors in germline-rescue models, whereas liver-specific *G6pc1* KO mice that retain measurable residual activity due to imperfect recombination develop HCA/HCC at high frequency (*30,32–33*). We therefore favor a model in which GSD1a-associated tumorigenesis requires convergence of cell-intrinsic glycogen storage with liver-wide metabolic dysfunction and systemic stress. One possible, untested contributor is the endocrine response to fasting hypoglycemia, including increased glucagon signaling, which could create a regenerative or growth-permissive environment in glycogen-loaded hepatocytes (*34-35*). This hypothesis is consistent with the clinical observation that GSD1a and GSD1b, both of which cause hypoglycemia in addition to glycogen accumulation, are associated with hepatic tumorigenesis in the absence of cirrhosis, whereas tumors in other glycogen storage diseases more often arise in the context of fibrosis or cirrhosis (*24*).

More broadly, our findings show that the physiological importance of metabolic zonation cannot be inferred solely from where an enzyme is most highly expressed or active. G6PC1-dependent glucose production is concentrated in periportal hepatocytes, yet *G6pc1* deletion from this zone was not sufficient to cause systemic metabolic manifestations of GSD1a. This result raises the possibility that, at least under standard laboratory conditions, zonation is not strictly required for basal systemic glucose homeostasis. However, our study does not determine whether maintaining homeostasis after zonal *G6pc1* loss carries a physiological cost. Defining that cost will require testing whether cross-zone compensation preserves systemic metabolism at the expense of metabolic efficiency, reserve capacity, or stress resilience.

Together, our findings separate local metabolic disruption from systemic metabolic disease and tumorigenesis. *G6pc1*-deficient hepatocytes in each targeted liver region accumulated glycogen, but this local storage phenotype did not progress to fasting hypoglycemia, lactate elevation, or liver tumor formation. These results support a model in which hepatic zonation operates within an organ-level buffering system that can absorb local metabolic defects before they progress to systemic disease.

## Supplemental Figures

**Figure S1.**
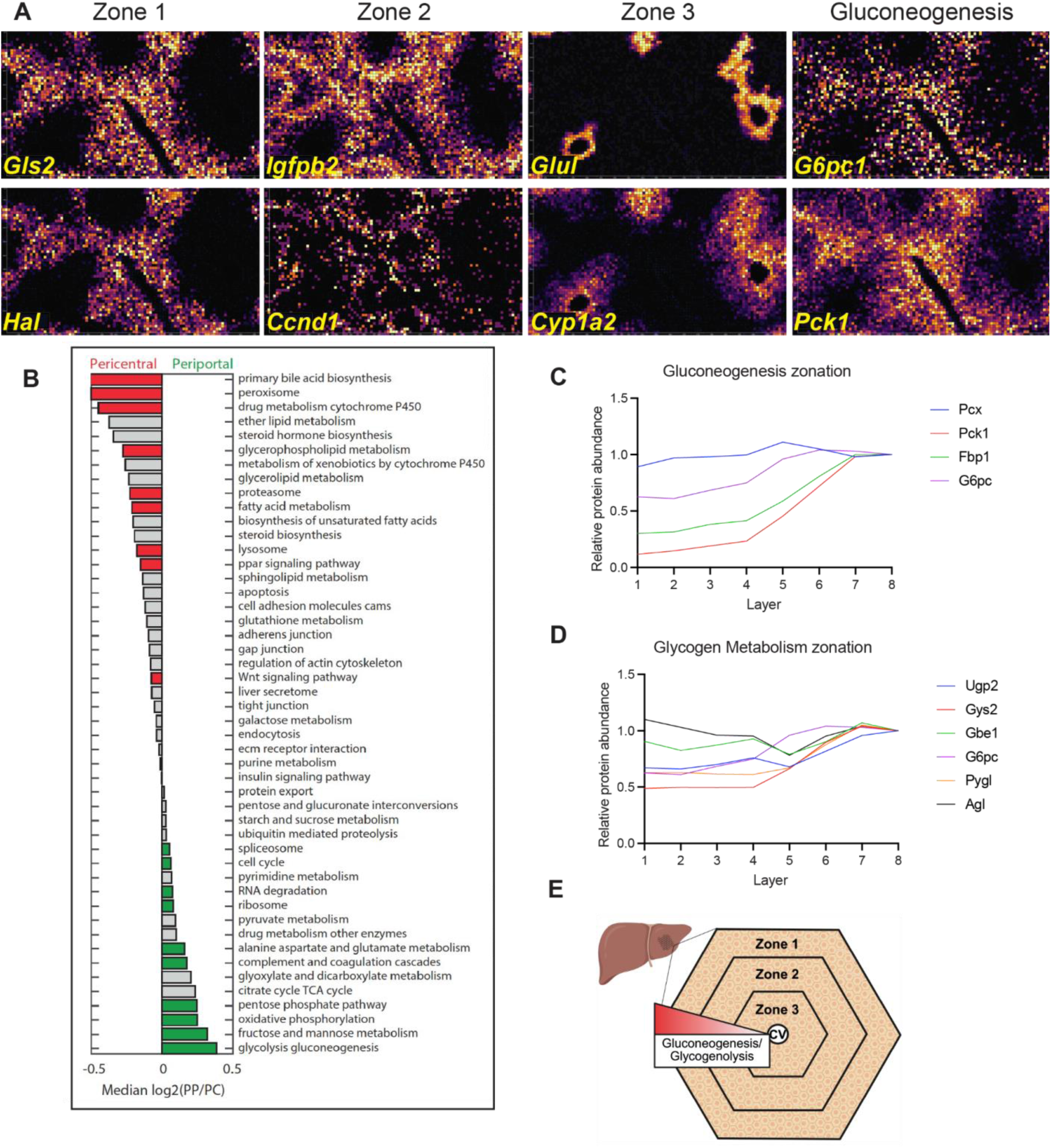
*G6pc1* and glycogen metabolism are enriched in periportal zone 1 hepatocytes. (**A**) Spatial transcriptomics (Xenium) showing *G6pc1* expression colocalizes with zone 1 markers as well as *Pck1,* another gene important for gluconeogenesis. (**B**) Reanalysis of published spatial proteomics data showing glycolysis/gluconeogenesis enrichment in zone 1 (*6*). Protein-level gradients for (**C**) gluconeogenesis and (**D**) glycogen metabolism enzymes from central vein (layer 1) to portal vein (layer 8). (**E**) Schematic model summarizing the periportal enrichment of G6PC1-dependent glucose production and glycogen metabolism.

**Figure S2.**
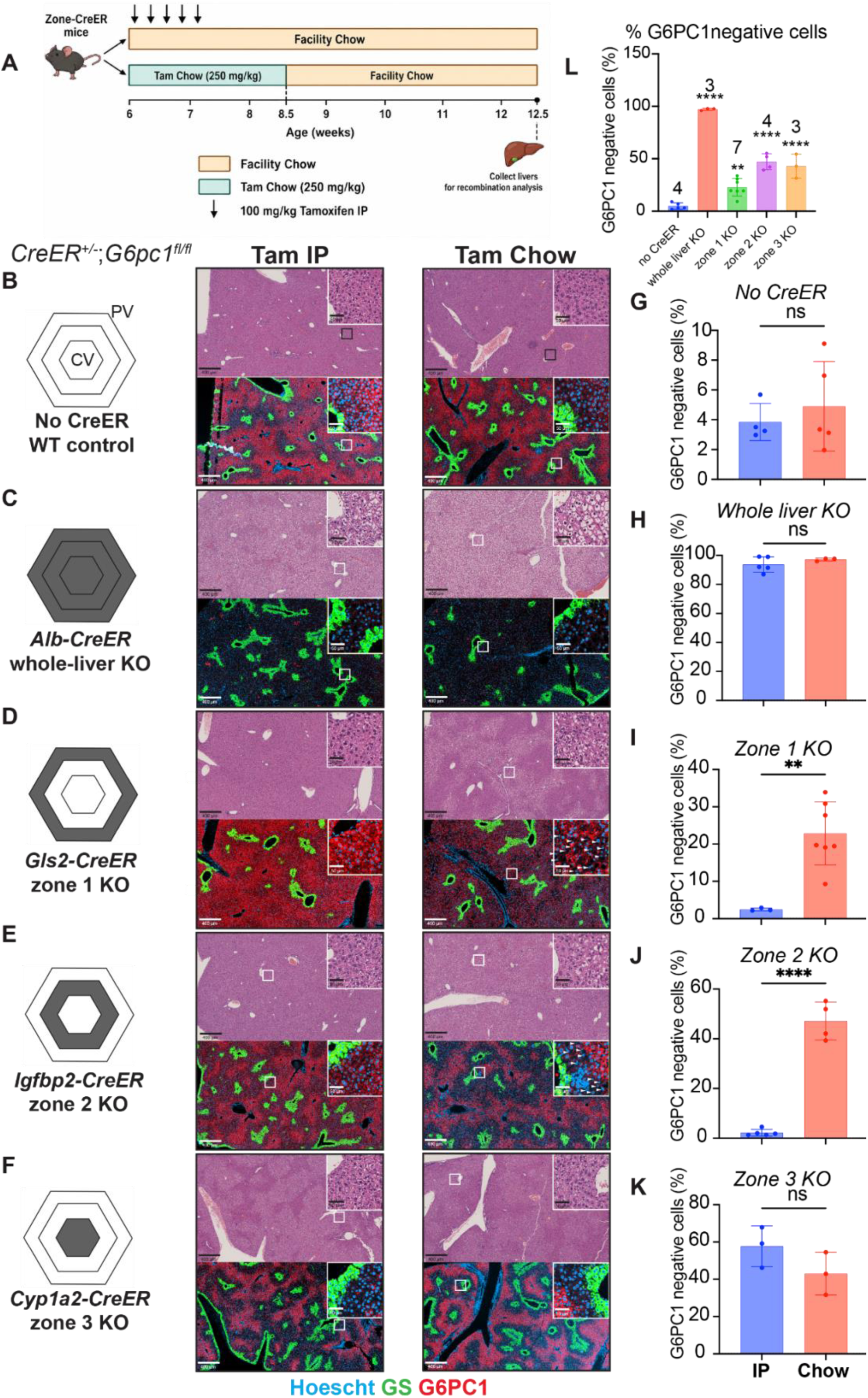
Tamoxifen chow improves recombination in zone-specific CreER models. (**A**) Experimental design comparing five consecutive daily IP tamoxifen injections with 2.5 weeks of tamoxifen chow for induction of *G6pc1* deletion. Mice were analyzed 1 month after completion of tamoxifen administration. (**B–F**) Representative H&E and G6PC1 immunofluorescence images from (**B**) no-CreER control, (**C**) whole-liver KO, (**D**) zone 1 KO, (**E**) zone 2 KO, and (**F**) zone 3 KO livers after IP tamoxifen or tamoxifen chow induction. GS staining marks pericentral hepatocytes and was used to orient the lobule. Arrows in (**D**) and (**E**) show G6PC1 negative cells seen with tamoxifen chow but not IP injection. (**G–K**) Quantification of G6PC1-negative cells as a percentage of all cells in the liver, after IP tamoxifen or tamoxifen chow induction in the corresponding models. Data represent mean ± SD. Unpaired two-tailed t tests were used to compare IP tamoxifen with tamoxifen chow within each model. Statistical significance was denoted as follows: *, P < 0.05; **, P < 0.01; ***, P < 0.001; ****, P < 0.0001.

**Figure S3.**
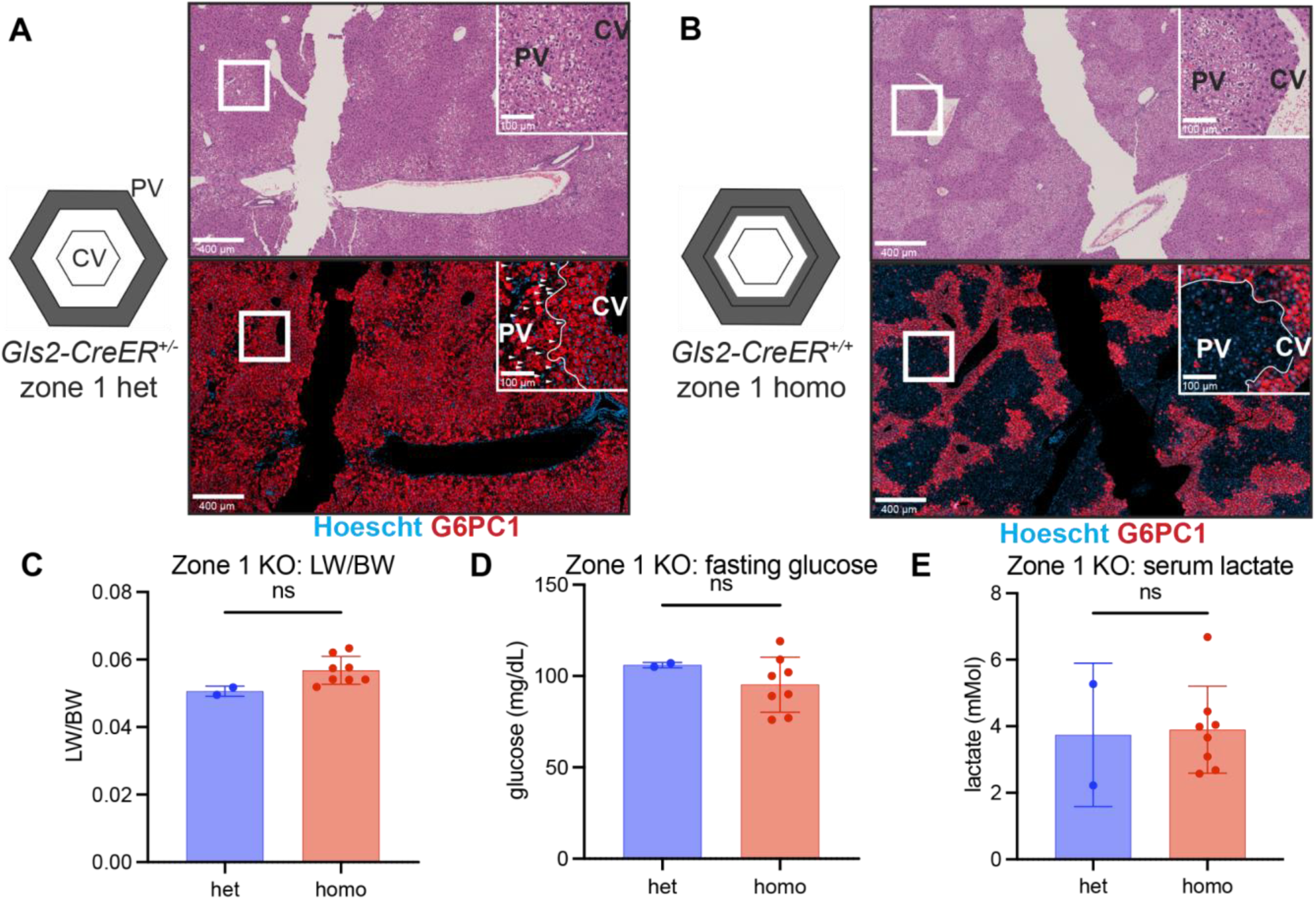
Increased zone 1 *G6pc1* recombination does not cause systemic GSD1a-like metabolic dysfunction. Representative H&E and G6PC1 immunofluorescence images from (**A**) *Gls2-CreER^+/−^;G6pc^fl/fl^*and (**B**) *Gls2-CreER^+/+^;G6pc1^fl/fl^* zone 1 KO livers 1 month after tamoxifen chow induction. White outlined regions indicate areas of G6PC1 loss used to identify recombined/KO regions. (**C**) Liver weight/body weight ratio, (**D**) fasting blood glucose, and (**E**) serum lactate in *Gls2-CreER^+/−^* and *Gls2-CreER^+/+^*zone 1 KO mice. Data represent mean ± SD. Unpaired two-tailed t tests were used to compare heterozygous and homozygous *Gls2-CreER* groups. Statistical significance was denoted as follows: *, P < 0.05; **, P < 0.01; ***, P < 0.001; ****, P < 0.0001.

**Figure S4.**
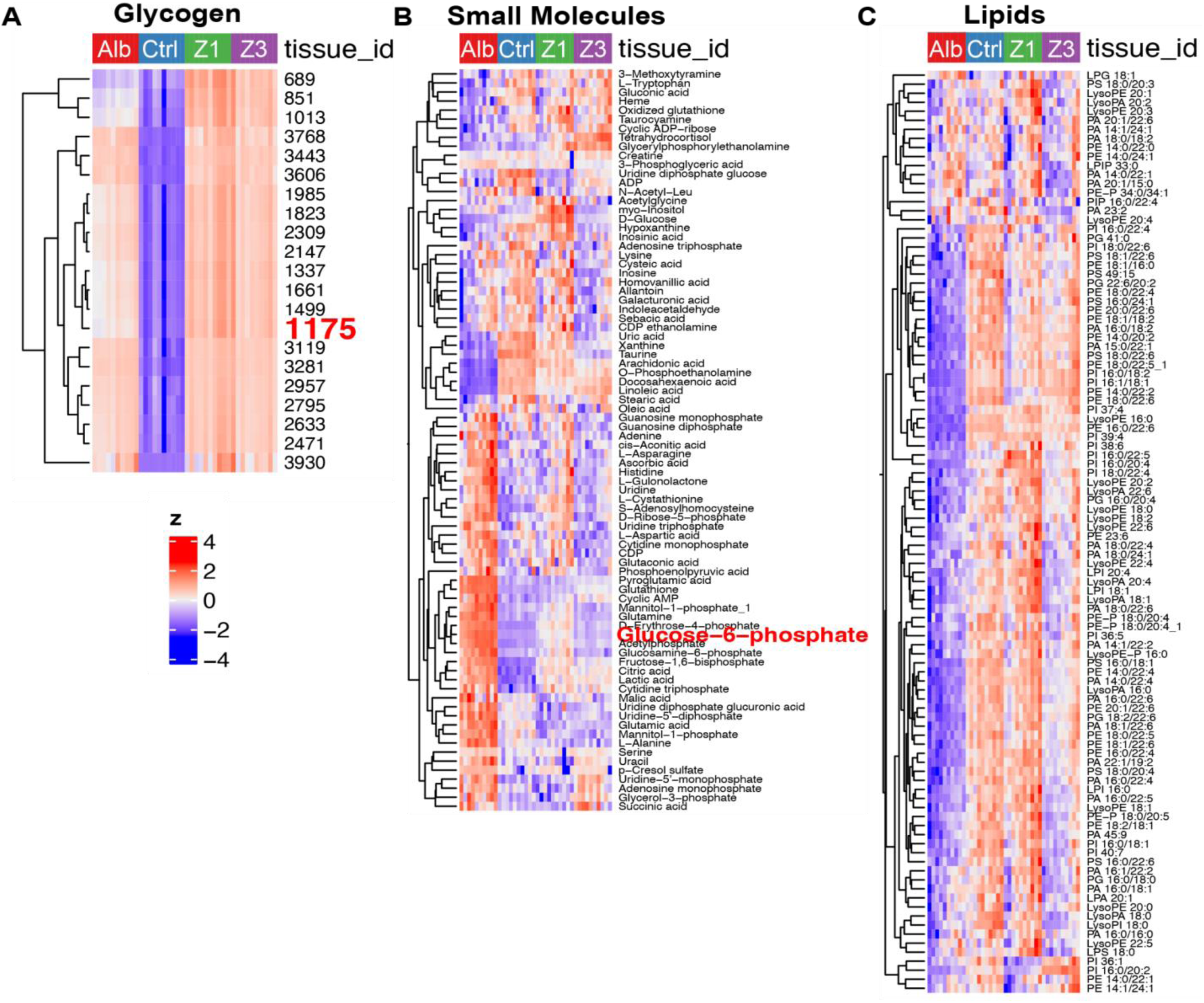
Heatmap of MALDI-ToF features in control, zonal *G6pc1* KO, and whole-liver KO liver sections. Heatmaps showing tile-level MALDI-ToF intensities for (**A**) glycogen-associated features, (**B**) aqueous small-molecule metabolites, and (**C**) lipid features in no-CreER control (Ctrl), zone 1 KO (Z1), zone 3 KO (Z3), and whole-liver KO (Alb) liver sections. Columns represent sampled spatial tiles from each tissue section, and rows represent detected features or annotated metabolites. Feature intensities are shown as row-scaled z scores. The glycogen-associated heatmap includes the representative glycogen feature m/z 1175, which is shown in Fig. 3A. The small-molecule heatmap includes glucose-6-phosphate, which is shown in Fig. 3B.

**Figure S5.**
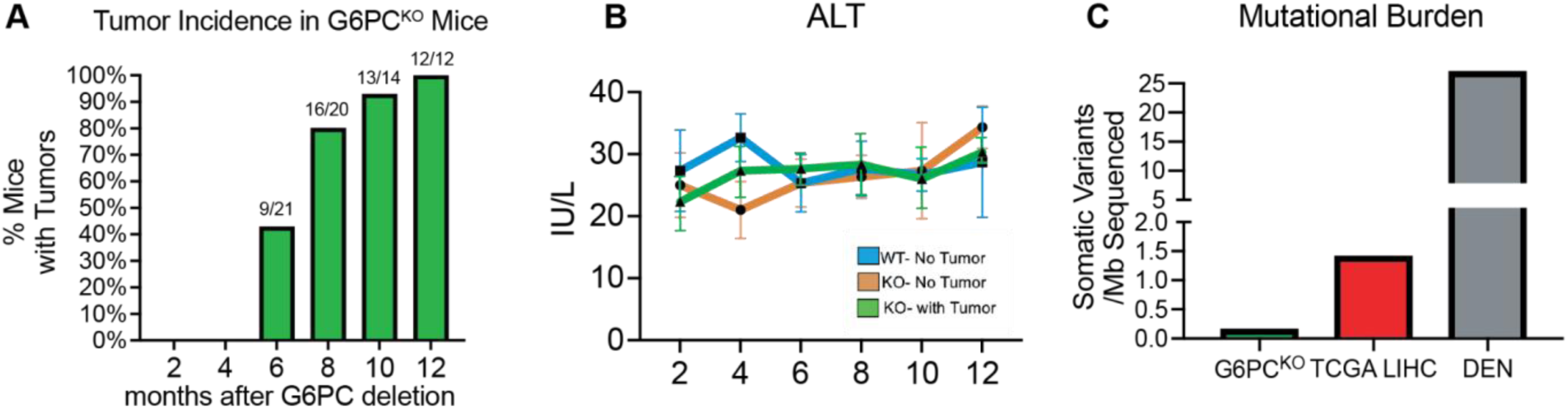
GSD1a tumors are not preceded by cirrhosis and have low mutational burden. (**A**) Over the course of 12 months, mice with *G6pc1* KO develop tumors that are not preceded by elevated serum (**B**) ALT. (**C**) These tumors have low mutational burden when compared to TCGA Liver Hepatocellular Carcinoma database or DEN-treated mouse livers.

## Materials and Methods

### Mouse strains and animal studies

*G6pc1^fl/fl^* mice were provided by Dr. Janice Chou at the National Institutes of Health. *Alb-CreER* mice were provided by Dr. Pierre Chambon at the University of Strasbourg. *Gls2-CreER* and *Cyp1a2-CreER* mice were established as previously described by Wei et al., and *Igfbp2-CreER* mice were established as previously described by Lin et al (*17*, *18*). All mice were maintained on a C57BL/6J background. To generate whole-liver or zone-specific *G6pc1* KO models, *G6pc1^fl/fl^* mice were crossed with *Alb-CreER*, *Gls2-CreER*, *Igfbp2-CreER*, or *Cyp1a2-CreER* mice. *G6pc1^fl/fl^* mice lacking *CreER* were used as genotype-matched colony controls. Male and female mice were used unless otherwise indicated, with sex indicated in individual experiments by closed circles for male mice and open circles for female mice. Unless otherwise indicated, mice were fasted for at least 9 hours prior to sacrifice, with fasting duration specified in the corresponding figure legends. For the 15-month aging cohort, mice were fasted for 4 hours prior to analysis. Fasting was initiated at noon, and mice did not have access to water during the fasting period. At the time of euthanasia, blood was collected by buccal bleed. Whole blood was then allowed to coagulate for 30 minutes at room temperature, followed by centrifugation at 2,000 x g for 10 minutes to isolate serum. For short-term experiments, mice were induced at 8 weeks of age and analyzed 1 month after completion of tamoxifen chow. Mice were housed in a specific pathogen-free facility under a 12-hour light/12-hour dark cycle wit h ad libitum access to food and water.

### Tamoxifen administration

For IP induction, tamoxifen was dissolved in corn oil at 10 mg/ml, and mice received 1 mg tamoxifen per mouse by IP injection once daily for 5 consecutive days. Each injection consisted of 100 μl tamoxifen solution. For chow-based induction, mice were fed tamoxifen-containing chow (250 mg/kg w/w; Envigo TD.130855) ad libitum for 2.5 weeks. For 1-month analyses, tissues and metabolic measurements were collected 1 month after finishing tamoxifen chow or IP injection regimens. All long-term zonal KO cohorts were induced using tamoxifen chow.

### AAV administration

To generate stochastic hepatocyte recombination, 8-week-old male *G6pc1^fl/fl^*mice lacking *CreER* were administered *AAV8-TBG-Cre* (Addgene 107787-AAV8) by retro-orbital injection. Mice received 2 × 10^9^, 1 × 10^10^, or 5 × 10^10^ vector genomes per mouse and were analyzed 1 month after injection.

### Blood and serum metabolite measurements

Blood glucose was measured immediately after sample collection using a Contour Next EZ glucometer (Contour Diabetes). Serum lactate was measured using the Megazyme L-Lactic Acid Assay Kit (Megazyme, catalog no. 700004310) according to the manufacturer’s instructions using a plate-based assay. Serum alanine aminotransferase (ALT) was measured at the UT Southwestern Metabolic Phenotyping Core using an Ortho Clinical Diagnostics Vitros 350 dry chemistry analyzer with reagent kit #1655281 (Ortho Clinical Diagnostics, Raritan, NJ), according to the manufacturer’s protocol. Additional serum analytes including triglycerides and uric acid were measured in selected experiments involving whole-liver KO and control mice (not shown).

### Histology and immunofluorescence

For routine histology, liver tissues were fixed in formalin for 48 hours, paraffin embedded, and sectioned at 4 μm thickness. Hematoxylin and eosin (H&E) staining was performed using standard protocols. For G6PC1 IF in Figures 1, 2, and 5, paraffin sections were used. For glycogen sensor studies (Figure 4), frozen liver sections were prepared by fixing tissues in formalin for 48 hours, followed by incubation in 30% sucrose for 24 hours. Tissues were then embedded in OCT compound at −20°C and sectioned at 10 μm using a Leica cryostat. Primary antibodies included anti-G6PC1 (Sigma-Aldrich, HPA052324; rabbit polyclonal, 1:100 for IF, diluted in 5% BSA in PBST) and anti-glutamine synthetase (BD Biosciences, #610518; mouse monoclonal, 1:500 in 5% BSA in PBST). Secondary antibodies included goat anti-mouse IgG1 Alexa Fluor 594 (Life Technologies, A-21125; 1:500) and donkey anti-rabbit IgG (H+L) DyLight 755 (Life Technologies, SA510043; 1:500). Nuclei were visualized using DAPI (Thermo Fisher Scientific, #5117; 1:1000). Images were acquired using either an Olympus IX83 fluorescence microscope or an Axioscan 7 slide scanner (Zeiss) through the UT Southwestern Whole Brain Microscopy Facility. Quantification of G6PC1 recombination was performed using QuPath software (version 0.5.0).

### Image quantification

Image analysis was performed on randomly generated annotations within selected regions of each liver section. Regions were manually adjusted only when necessary to exclude tissue folds, sectioning artifacts, or poorly preserved tissue. Cells were identified using nuclei, and cytoplasmic regions surrounding each nucleus were used to quantify G6PC1 immunofluorescence and glycogen detector signal. For Figures 1, 2, and 5, G6PC1 recombination was estimated by thresholding G6PC1 immunofluorescence across nuclei-associated cellular regions. Because non-parenchymal cells were not excluded from this analysis, the percentage of G6PC1-negative cells represents an image-based estimate of tissue-level G6PC1 loss rather than a hepatocyte-specific recombination frequency. For Figure 4, cells with nuclear area <50 µm² were excluded to enrich for hepatocytes and minimize the contribution of non-parenchymal cells. A more stringent G6PC1 intensity threshold was then applied to classify hepatocytes as G6PC1-high or G6PC1-low based on cytoplasmic G6PC1 signal. We used the term G6PC1-low, rather than G6PC1-negative, because this threshold was designed to capture both recombined hepatocytes in *G6pc1* knockout livers and the physiologic low-G6PC1 hepatocyte population present in control livers, allowing these populations to be compared directly. Glycogen abundance was estimated from the mean cytoplasmic fluorescence intensity of the glycogen detector.

### Xenium spatial transcriptomics

Previously generated formalin-fixed paraffin-embedded (FFPE) Xenium spatial transcriptomic data from mouse liver were reanalyzed from our prior study (*8*). In that study, fresh livers were harvested, and the largest lobe was selected for fixation. Peripheral regions were trimmed to facilitate formalin penetration, and tissues were fixed in 10% neutral-buffered formalin for 24 hours at room temperature before transfer to 70% ethanol. Samples were paraffin embedded, sectioned at 4 μm thickness, and submitted to BioChain Institute Inc. for spatial transcriptomic profiling using the 10x Genomics Xenium platform. A custom Xenium panel targeting 440 genes enriched for liver zonation and metabolic pathways was used. Raw Xenium-formatted spatial transcriptomic data were processed using the spatialdata-io module (v0.1.5). Data were imported using the Xenium function and underwent quality control filtering to remove cells with fewer than 20 total transcript counts and genes expressed in fewer than 5 cells. Filtered data were normalized using scanpy.pp.normalize_total (Scanpy v1.10.3), followed by logarithmic transformation with <u>scanpy.pp</u>.log1p. Principal component analysis (PCA) was performed on normalized data. The top 50 principal components were evaluated using variance ratio plots, and the first 20 principal components were selected for downstream analyses including neighborhood graph construction and UMAP embedding (min_dist = 0.5, spread = 1.0, random_state = 0). Clustering was performed using the Leiden algorithm across multiple resolutions, and resolution 1.0 was selected on the basis of cluster interpretability and marker expression. Clusters were annotated by hepatic zone and cell type using established marker genes, including zone-specific hepatocyte markers such as *Gls2* and *Cyp1a2*, as well as markers for bile duct cells, Kupffer cells, endothelial cells, hepatic stellate cells, and lymphocytes. Xenium data were used primarily to confirm the periportal enrichment of gluconeogenic gene expression.

### Reanalysis of published spatial proteomics

Previously published spatial proteomics data from Ben-Moshe et al. (2019) were used to examine the zonal distribution of proteins involved in gluconeogenesis and glycogen metabolism (*6*). Rather than reprocessing raw proteomic data, published layer-resolved protein abundance plots were used to evaluate gradients across the porto-central axis. Proteins involved in gluconeogenesis and glycogen metabolism, including G6PC1, glycogen synthase (GYS2), glycogen branching enzyme (GBE1), and other periportal-enriched enzymes, were examined for spatial enrichment patterns. Published KEGG pathway analyses demonstrating periportal enrichment of glycolysis/gluconeogenesis were adapted to illustrate the strong zonation of this metabolic pathway. Figures were generated by selective representation of published layer-specific enrichment patterns to highlight proteins relevant to hepatic glucose production.

### Spatial metabolomics imaging and analysis

At the time of collection, livers were placed in weighboats positioned on liquid nitrogen for rapid freezing. Frozen tissues were sectioned at 10 μm thickness using a Leica CM1860 cryostat at −20°C. Sections were mounted onto Indium Tin Oxide (ITO) IntelliSlides (Bruker, ref. #1868957) for metabolomic and lipidomic imaging and onto Electron Microscopy Sciences slides (EMS, cat. no. 71873-02) for glycomic imaging. Slides were desiccated under vacuum for at least 1 hour prior to matrix application. For metabolomic and lipidomic imaging, N-(1-naphthyl)ethylenediamine dihydrochloride (NEDC) matrix was applied uniformly using an HTX M5 pneumatic sprayer (14 passes, 0.06 mL/min flow rate, 3 mm offset, 1200 mm/min velocity, 30°C nozzle temperature, 10 psi nitrogen pressure, 50°C heated tray). For glycomic and glycogen imaging, EMS slides underwent antigen retrieval followed by enzymatic treatment with isoamylase and PNGase, after which alpha-cyano-4-hydroxycinnamic acid (CHCA) matrix was applied using the same sprayer (10 passes, 0.100 mL/min flow rate, 3 mm offset, 1300 mm/min velocity, 79°C nozzle temperature, 10 psi nitrogen pressure, 50°C heated tray). Slides were scanned to generate TIFF images and analyzed using a Bruker timsTOF flex MALDI imaging instrument. For small-molecule and lipid detection, acquisition parameters included 65% laser power, 396 laser shots per pixel at 10,000 Hz, a 46 μm × 46 μm laser raster for 50 μm × 50 μm pixels, negative polarity, scan range of 20–1400 m/z, and external calibration using tune mix solution immediately prior to acquisition. For glycogen detection, acquisition parameters included 37% laser power, positive polarity, scan range of 500–4000 m/z, and optimized settings for high-mass glycogen chain detection. Raw datasets were processed using SCiLS Lab (Bruker Daltonics) with spectral alignment and normalization to total ion count (TIC). Peak annotation was manually performed against an in-house curated spectral library, and annotated peak lists were exported as .csv files containing per-pixel ion intensities. Spatial metabolomics analyses were performed in R using processed ion-intensity matrices and exclusion masks. For glycogen analysis, glycogen features were analyzed together with macroscopic exclusion masks generated from the combined glycan and glycogen feature space. The glycogen ion at m/z 1175 was visualized after log10(x + 1) transformation with masked pixels excluded. For small-molecule and lipid analyses, TIC-normalized matrices were further processed using G6P-based exclusion masks and PCA-based inclusion masks. G6P was visualized using percentile-trimmed color scaling. For pseudobulk analyses, pixels were grouped into fixed 500 × 500 tiles after exclusion of a 50-unit tissue-edge margin. Tiles were retained if they contained at least 50 pixels and less than 5% masked area. For glycogen datasets, mean log2(x + 1) intensity was calculated per tile; for small-molecule and lipid datasets, median intensity was calculated per tile. Tile-level matrices were centered and scaled prior to principal component analysis. Heatmaps were generated using 10 randomly sampled tiles per tissue after per-feature z-scoring.

### Glycogen sensor design, purification, and staining

A fluorescent glycogen sensor was generated using a recombinant mCherry-6xHis-FLAG-CBM20 fusion protein cloned into a pET bacterial expression vector (VectorBuilder). The glycogen-binding domain consisted of residues 261–358 of human STBD1, corresponding to the CBM20 glycogen-binding domain. The construct was expressed in Escherichia coli BL21(DE3) cells (New England Biolabs, #C2527). Individual colonies were selected and cultured in 5 mL terrific broth (TB), then transferred to 1 L TB after 16 hours of rocking incubation at 30°C. Recombinant protein was purified using Ni-NTA affinity purification and stored in a buffer containing 50 mM Tris (pH 7.5), 50 mM NaCl, 10 mM MgCl₂, 30% glycerol, and 50 mM KCl. For glycogen staining, liver tissues were fixed in 10% formalin for at least 24 hours, transferred to 30% sucrose for at least 24 hours, embedded in OCT, frozen, and cryosectioned at 10 μm thickness. Sections were allowed to dry for approximately 15 minutes before outlining with a hydrophobic barrier pen. Slides were washed with PBS containing 0.25% Triton X-100 (PBST) three times for 2 minutes each. For specificity controls, sections were incubated with either 1% α-amylase in PBS or PBS alone for 30 minutes at room temperature. Slides were then incubated in 5% BSA in PBST containing glycogen sensor (1:1000) and Hoechst (1:1000) for 30 minutes at room temperature, followed by three PBST washes and mounting with AquaMount for imaging. For multiplex staining with G6PC1 IF, frozen sections were first rehydrated through a xylene-to-water gradient followed by antigen retrieval for 20 minutes in Antigen Retrieval Citra Plus buffer (BioGenex, #HK080) at sub-boiling temperature and allowed to cool to room temperature. After washing and blocking in 5% BSA in PBST for 30 minutes, primary antibody against G6PC1 (1:100) and glycogen sensor (1:1000) were applied simultaneously overnight. The following day, sections were washed and incubated with far-red anti-rabbit secondary antibody (Invitrogen, SA5-10043, 1:500) and Hoechst (1:1000) in 5% BSA in PBST, followed by washing, mounting, and imaging. All glycogen sensor experiments were performed using frozen sections.

### Tumor studies and long-term aging cohorts

For long-term tumor studies, mice were induced with tamoxifen chow and aged for 10, 12, or 15 months following *G6pc1* deletion. Both male and female mice were included, and exact cohort sizes are indicated in the corresponding figures. Mice were monitored until scheduled euthanasia and were not removed early for morbidity. Tumor burden was assessed qualitatively by gross liver examination and by calculation of liver weight-to-body weight (LW/BW) ratios. Representative livers from each genotype were additionally evaluated by hematoxylin and eosin staining for histologic assessment of nodules and tissue architecture. Histologic evaluation was performed without formal veterinary pathology review.

### Whole-exome sequencing analysis

Genomic DNA was isolated from transformed and non-transformed *G6pc1* KO liver tissue using the QIAamp DNA Blood Midi Kit (Qiagen). DNA concentration and quality were assessed using a NanoDrop spectrophotometer (ND-100). Library preparation was performed using the SureSelect Mouse Kit (Agilent) according to the SureSelectXT Library Kit and SureSelectXT Target Enrichment System for Illumina (version B.2). Sequencing was performed by Psomagen using an Illumina HiSeq 2500 platform with 150-bp paired-end reads to achieve approximately 200× raw sequencing depth. Somatic variants were called using a cutoff of 0.1 allele frequency in tumor samples and were excluded if present in dbSNP (https://www.ncbi.nlm.nih.gov/snp/). Tumor mutational burden was calculated as somatic variants per megabase and compared with publicly available TCGA liver hepatocellular carcinoma (LIHC) datasets and published diethylnitrosamine (DEN)-induced mouse liver tumor datasets.

## Statistical analysis

Data are presented as mean ± SD unless otherwise indicated. No statistical method was used to predetermine sample size. All biological replicates represent individual mice (n = 1 mouse). For MALDI-ToF analyses, spatial tiles were used for visualization, heatmap generation, and dimensionality reduction. Because tiles from the same tissue section are spatially correlated and do not represent independent biological replicates, genotype-level statistical inference was not performed from tile counts. Statistical analyses were performed using GraphPad Prism version 10 and R. Standard parametric tests were used throughout the study. Comparisons among multiple groups were performed using one-way or two-way ANOVA as indicated in the figure legends. One-way ANOVA analyses were followed by Tukey’s multiple comparisons test. Two-way ANOVA analyses were followed by Fisher’s least significant difference (LSD) test for multiple comparisons. Statistical significance was defined as P < 0.05. In figures, significance was denoted as follows: *, P < 0.05; **, P < 0.01; ***, P < 0.001; ****, P < 0.0001.

## Study approval

All animal studies were performed in accordance with protocols approved by the Institutional Animal Care and Use Committee (IACUC) at UT Southwestern Medical Center under Animal Protocol 2022-102897. No human samples were used in this study.

## Acknowledgments

We thank members of the Zhu and DeBerardinis laboratories for helpful discussions, technical advice, and feedback throughout this project. We thank Dr. Denise Ramirez from the UT Southwestern Whole Brain Microscopy Facility for imaging support, the UT Southwestern Metabolic Phenotyping Core for serum chemistry measurements, and the UT Southwestern Tissue Management Shared Resource for histology support. Additionally, we thank Lauren Zacharias and Dr. Thomas Matthews from the Children’s Research Institute Metabolomics Facility for metabolomics assistance.

## Author contributions

Conceptualization: R.L., H.Z., and R.J.D. Methodology: R.L., L.C., P.T.N., R.R., A.R., R.C.S., H.Z., and R.J.D. Investigation: R.L., L.C., P.T.N., S.K., M.B., T.T., E.C., R.R., and A.R. Formal analysis: R.L., S.K., R.R., A.R., and R.C.S. Resources: J.C., R.C.S., H.Z., and R.J.D. Writing – original draft: R.L. Writing – review and editing: R.L., L.C., P.T.N., S.K., M.B., T.T., E.C., R.R., A.R., J.C., R.C.S., H.Z., and R.J.D. Supervision: R.C.S., H.Z., and R.J.D. Funding acquisition: R.L., P.T.N., R.C.S., H.Z., and R.J.D.

## Data and materials availability

Data needed to evaluate the conclusions in this study are present in the paper and/or the Supplementary Materials. Whole-exome sequencing and MALDI-ToF spatial metabolomics data will be deposited in public repositories prior to publication; accession numbers are pending. Processed image quantification data and source data for quantified figures will be available from the corresponding authors upon reasonable request. The *G6pc1^fl/fl^*mouse line was provided by J.C. and is available subject to institutional and material transfer agreements. The glycogen sensor construct and other materials generated in this study are available from the corresponding authors upon request.

## Supplementary Materials

This PDF file includes:

Figs. S1 to S5

Legends for Figs. S1 to S5

